# Nucleus Reuniens: Modulating Negative Overgeneralization in Periadolescents with Anxiety

**DOI:** 10.1101/2023.11.14.567068

**Authors:** M. Vanessa Rivera Núñez, Dana McMakin, Aaron T. Mattfeld

## Abstract

**Background:** Anxiety affects 4.4-million children in the United States with an onset between childhood and adolescence, a period marked by neural changes that impact emotions and memory. Negative overgeneralization – or responding similarly to innocuous events that share features with past aversive experiences – is common in anxiety but remains mechanistically underspecified. The nucleus reuniens (RE) has been considered a crucial candidate in the modulation of memory specificity. Our study investigated its activation and functional connectivity with the medial prefrontal cortex (mPFC) and hippocampus (HPC) as neurobiological mechanisms of negative overgeneralization in anxious youth.

**Methods:** As part of a secondary data analysis, we examined data from 34 participants between 9-14 years (mean age ± SD, 11.4 ± 2.0 years, 16 females) with varying degrees of anxiety severity. During the Study session participants rated images as negative, neutral, and positive. After 12-hours, participants returned for a Test session, where they performed a memory recognition test with repeated (targets) and similar (lures) images. Labeling negative relative to neutral lures as “old” (false alarms) was our operational definition of negative overgeneralization.

**Results:** Negative relative to neutral false alarmed stimuli displayed elevated RE activation (at Study and Test) and increased functional connectivity with the CA1 (at Test only). Elevated anxiety severity was associated with reductions in the RE-mPFC functional coupling for neutral relative to negative stimuli. Exploratory analyses revealed similar patterns in activation and functional connectivity with positive stimuli.

**Conclusions:** Our findings demonstrate the importance of the RE in the overgeneralization of memories in anxious youth.

## INTRODUCTION

The onset of anxiety coincides with the period between childhood and adolescence, a developmental window marked by neurobiological upheaval (1–6). Negative overgeneralization – responding similarly to innocuous events that share attributes with past aversive memories – is a common anxiety feature (7–9) that takes root during this period and is mechanistically poorly understood. The nucleus reuniens (RE) of the ventral midline thalamus – well-connected to memory related regions (10–15) – may be an important contributor to negative overgeneralization. Identifying neural mechanisms subserving negative overgeneralization can play an important role in understanding the course of anxiety disorders (16–18). For instance, in generalized anxiety disorder (GAD), it is not uncommon for the emotional response to a negative event (e.g., being bullied at school) to transfer to seemingly related non-threatening scenarios (e.g., fear of making friends at the park) (8). The persistence of these negative expectations can generalize across contexts, leading to disruptive symptoms (e.g., worry and avoidance) that could give rise to significant impairment (19–22).

The amygdala has been a central focus of mechanistic investigations in negative overgeneralization (23–29). In non-human models, lateral amygdala neurons shift from cue-specific to generalizing responses following conditioning (24), while basolateral amygdala neurons exhibit shallow auditory tuning curves for perceptually similar unconditioned stimuli (CS-) (23). Similarly, in humans, elevated amygdala activation during fear conditioning correlated with the subsequent generalization of aversive images (27–29).

Despite the amygdala’s established contribution to memory generalization, regions such as the medial prefrontal cortex (mPFC), hippocampus (HPC), and ventral midline thalamic nuclei offer compelling alternative targets to investigate within the context of negative overgeneralization (8,9,25,27,30–35). The mPFC shares direct and indirect connections with the amygdala and HPC (36) and is thought to exercise modulatory control of emotional and mnemonic processes that support fear generalization (33,36,37). The HPC consists of unique subfields that have distinct functional properties that support learning and memory (e.g., pattern separation and completion) (38–40). The midline thalamus, comprised of the paraventricular (PV), paratenial (PT), RE and rhomboid (RH) nuclei (12,41,42), has a wide array of cortical and subcortical inputs from learning and memory structures (12). However, unlike the PV and PT nuclei, the RE and RH have overlapping projections to the mPFC and HPC, with the RE having comparatively stronger connectivity, making it critical memory processing (14,43–46). The aforementioned areas are mechanistically important in learning (14,32) and implicated in disorders related to deficits in memory and emotion regulation (14,47,48). However, their mechanistic contributions may only tell part of the story in the context of development.

The transition between childhood and adolescence is characterized by dynamic neurobiological changes (29,49–53). In the HPC, rapid maturation of the CA1 compared to the CA3/DG (49) has been proposed to favor memory overgeneralization in children with anxiety (29). The protracted synaptic pruning and underlying myelination of the prefrontal cortex (50–52,54) may contribute to alterations in the role of this region in mnemonic discrimination (29). In the case of the midline thalamus, the only study that explored its development (53) showed the significant overlap across all midline thalamic nuclei in infant compared to adult rats, as well as contralateral projections to cortical sites that became ipsilateral with age. Taken together, maturational changes between childhood and adolescence highlight the importance of investigating the functional interaction of the amygdala, mPFC, HPC, and RE during this developmental window, especially in the context of memory generalization and anxiety.

The current study is a secondary analysis of a project that examined the neural correlates of negative overgeneralization in anxious youth (29). Here, we aim to investigate the role of the RE in negative overgeneralization. We specifically predicted that activation of the RE would increase for negative relative to neutral stimuli that were false alarmed (FA). Based on the connectivity profile of the RE, we hypothesized that the functional connectivity of RE with the CA1 and mPFC would increase for negative relative to neutral FA items. To examine anxiety as a moderator in the activation and functional connectivity of the RE, we predicted that elevated anxiety severity would lead to greater differences between negative relative to neutral FA images.

## METHODS and MATERIALS

### Participants

Fifty-two participants were enrolled in the study, however, the final sample consisted of 34 youth (mean age ± SD, 11.4 ± 2.0 years, 16 females). Eighteen participants were not included due to: withdrawing from the study [1], left-handedness [3], experimenter errors in administering the task [5], did not show up for scheduled visit [2], excessive motion [1], and poor behavioral performance defined as target hit rates of 1.5SD below average performance [6]. Florida International University’s local institutional review board approved our study procedures and methods. Parents and children provided informed consent and assent, respectively. Youth between the ages of 9 and 14 years participated in an emotional mnemonic similarity task while undergoing two different fMRI sessions. To enhance our effect of interest (i.e., negative overgeneralization), we selected the age range that captured the onset of prepuberty and recruited participants from community samples and a university clinic for anxious youth. Please see (29) for a detailed description of study design and procedures. Anxiety severity was assessed by means ofthe Pediatric Anxiety Rating Scale (PARS-6; (55)). Participants were excluded based on MR contraindications, comorbid or stand-alone diagnoses not related to anxiety and a Weschler Abbreviated Scale of Intelligence–Second Edition (WASI-II) full-scale IQ score (56) – below 80.

### Procedure

Participants completed a clinical intake session and were subsequently randomized to either a sleep (N = 16) or wake (N =18) condition. For the purposes of this paper, however, sleep versus wake differences on negative overgeneralization were beyond the scope of our research question. Therefore, we collapsed across conditions to increase our power and investigate the midline thalamic mechanisms underlying memory generalization in anxious youth. A week later, participants returned to start the emotional similarity task (EMST) (29,57) (Figure 1). During the Study session emotionally valenced images were presented in the center of the screen for 2 seconds. Participants rated image valence while on the screen using an MRI - compatible response device. After each picture a white fixation cross on a black screen appeared for 2-6 seconds. A total of 145 stimuli (48 negative, 47 neutral, 50 positive) were divided between two scan runs, each lasting about 8-minutes. During the Test session participants were shown targets (i.e., old images), foils (i.e., new images), and lures (i.e., similar images). The stimulus presentation timing and inter-stimulus-interval were the same across sessions.

**Figure 1.**
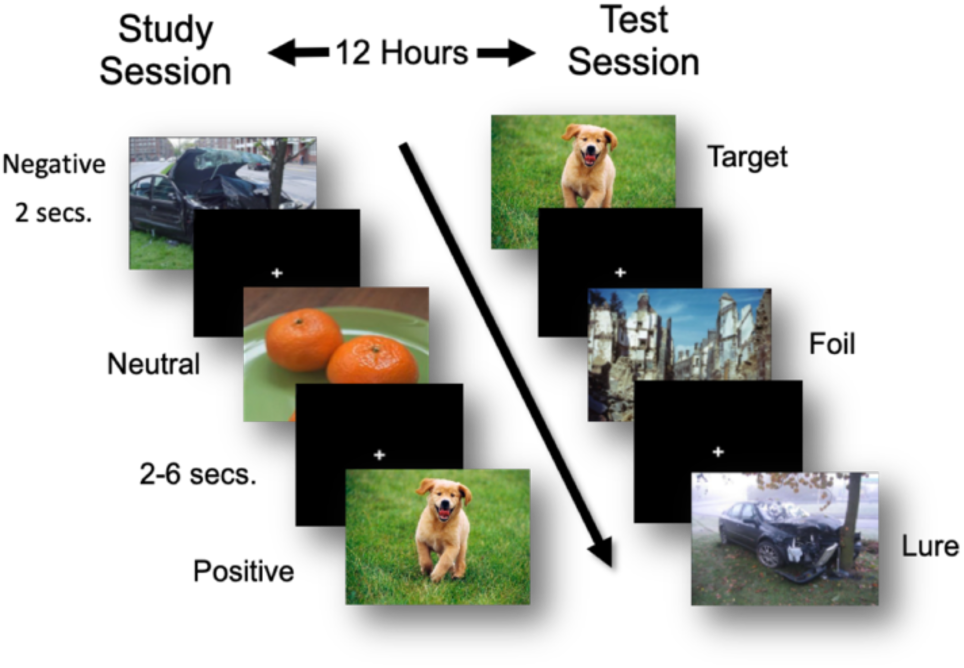
Emotional Similarity Task. The task began with a Study Session inside the scanner where participants were shown negative, neutral, and positive valenced images. Participants were instructed to provide valence ratings (i.e., negative, neutral, or positive) while the pictures remained on the screen for 2 seconds each. After 12 hours, all volunteers returned for the Test Session scan where they were shown targets (i.e., images presented during the first visit), foils (i.e., new images), and lures (i.e., images subtly different from the original ones). Participants were instructed to label the images as ‘old’ or ‘new’. If the picture was *exactly* the same as the one presented during the Study Session, the image was considered ‘old’. The duration of the stimulus presentation, as well as the interstimulus time was the same as the Study Session scan.

Participants indicated whether the images were ‘old’ or ‘new’ using an MRI-compatible response device. They were instructed to endorse ‘old’ only if the image was exactly the same as before. If participants responded to lures as ‘old,’ these trials were labeled false alarms (FA) and constituted our definition of generalization. Lures that were identified as ‘new’ were considered correct rejections (CR). Identifying targets as ‘old’ were considered hits (target hits), while targets labeled ‘new’ were considered misses (target misses). Foils labeled ‘old’ were considered FA, while those labeled ‘new’ were considered CR. During the Test session, a total of 284 stimuli were presented (16 negative targets, 16 neutral, and 16 positive targets, 32 negative lures, 33 neutral lures, 33 positive lures, 42 negative foils, 49 neutral foils, 48 positive foils), split between 4 runs with each lasting about 7 minutes.

### MRI Data Collection

Data were acquired using a Siemens 3T Magnetom Prisma scanner with a 32-channel head coil. A T1-weighted anatomical scan (i.e., TR/TE = 2500/2.9 ms, flip angle = 8°, voxel size = 1 mm isotropic, field of view = 256 mm, 176 sagittal slices), T2*-weighted functional scan (TR/TE = 993/30 ms, flip angle = 52°, voxel size = 2.4 mm isotropic, field of view = 216 mm, 56 axial slices, slice acceleration = 4; first 4 volumes were discarded), and diffusion-weighted images (DWI) (1.7 mm^3^, slice acceleration = 3, 96 directions, seven b = 0 frames, and four b-values [six 500s/mm2 directions, fifteen 1000s/mm2 directions, fifteen 2000s/mm2 directions, and sixty 3000s/mm2 directions]) were collected. For distortion correction purposes, we collected a fieldmap in opposite phase encode direction from the original DWI phase encode direction.

### fMRI Data Preprocessing

All neuroimaging data were preprocessed using a custom Nipype (Nipype version 0.12.1 (58) pipeline drawing from the following software: Analysis of Functional Neuroimages (AFNI version 16.3.18 (59)), FMRIB Software Library (FSL version 5.0.10 (60)), FreeSurfer (version 6.0.0 (61)), and ANTs. Functional data were motion corrected to the first volume of the first run utilizing FSL’s McFlirt (62). Motion and intensity outliers were identified (>1 mm frame-wise displacement; >3 SD mean intensity), then we spatially filtered our data using a 4 mm kernel using the SUSAN algorithm (63).

### Anatomical Regions of Interests

Freesurfer’s aparc+aseg.mgz file was utilized to create an anatomical region of interest mask of the mPFC, which included the lateral and medial orbitofrontal cortex, the rostral anterior cingulate cortex, and the superior frontal cortex. Hippocampal subfields were segmented according to a consensus labeling approach (29). Expertly labeled manual segmentations from an in-house atlas set of 19 young adults with both T1 MPRAGE (0.75 mm isotropic) and T2-FSE (0.47×0.47 mm2 in-plane, 2.0 mm thickness; oblique acquisition along longitudinal axis of hippocampus) scans were normalized to unlabeled subject T1-weighted images and join label fusion was applied (64). Following atlas normalization to the unlabeled subject, a weighted averaging technique was utilized for subfield labeling taking into account the unique label and intensity information contributed by each member of the atlas (29).

### Identification of the Nucleus Reuniens

We utilized probabilistic tractography combined with a data-driven k-means clustering approach (61) to approximate the location of the RE. For further details, please see Supplementary section: “Probabilistic Tractography and K-Means Clustering.”

### Functional Data Analysis

General linear models for the Study and Test sessions were used to examine the neural correlates of negative overgeneralization. Each model included the following nuisance regressors: motion (x, y, z translations and pitch, roll, and yaw rotations), first and second derivatives of motion, normalized motion, Lagrange polynomials (first through third order) to account for low frequency signal variability, and regressors for each time point exceeding our motion outlier thresholds. Regressors of interest included lure and foil FAs and CRs, and target hits and misses for the 3 valences (negative, neutral, and positive). The Study session model included a similar set of regressors that were derived from Test session behavior.

### Functional Connectivity Analysis

To assess the frontotemporal functional connectivity with the RE, we conducted a beta-series correlation analysis (65). Each trial was modeled using a least-square single approach (66). Contrast parameter estimates for negative and neutral lure FAs were combined into separate beta-series (i.e., negative and neutral) and the activation was averaged across voxels for each region of interest (i.e., mPFC, RE, CA1). Pearson’s correlation coefficients and their respective arctangents were calculated for the RE-CA1 and RE-mPFC correlations.

### Analytic approach

We first extracted contrast parameter estimates for the RE. To assess changes in the activation with the RE for negative relative to neutral lure images across Study and Test sessions, we conducted a mixed general linear model (MGLM). Valence (negative vs. neutral), Session (Study vs. Test), and Subjects were included in the model, with Subjects being treated as random effects. Anxiety severity (Pars-6 scores) was included in all models to assess its influence on the RE and memory generalization. For all analyses, if no significant interactions were identified, main effects of valence and session were assessed followed by post-hoc paired samples t-tests, comparing differences in activation or functional connectivity for negative relative to neutral lure images that were FA (Test session) or for those that were subsequently replaced by a lure and FA (Study session). A Shapiro-Wilk test was utilized to assess violations in normality. If a violation was identified, a non-parametric test was implemented (i.e., Wilcoxon signed rank test). To minimize Type I error, we conducted a Holm-Sidak multiple comparisons correction.

For our functional connectivity analyses we conducted two separate MGLM, one for the RE-CA1 and another for the RE-mPFC. Each model included Valence, Session, and Anxiety severity. Subjects were treated as random effects. As before, if no significant interactions were identified, we assessed for main effects of valence and session followed by post-hoc paired samples t-tests that compared functional connectivity differences between negative and neutral lure images that were FA (Test session) or for those that were subsequently replaced by a lure and FA (Study session). A Shapiro-Wilk test examined violations in normality. If a violation was identified, a Wilcoxon signed rank test was utilized. To minimize Type I error, we conducted a Holm-Sidak multiple comparisons correction.

## RESULTS

### The nucleus reuniens participates in the generalization of negative stimuli

To assess whether the RE contributed to negative overgeneralization, our MGLM first examined the influence of anxiety on activation of the RE across sessions. Our results did not reveal a significant 3-way interaction (Valence X Session X Pars-6: negative vs. neutral: z = −0.57, *p* = 0.57, 95% CI [-0.09, 0.05]; positive vs. neutral: z = −0.53, *p* = 0.60, 95% CI [-0.09, 0.05]).

Similarly, we did not observe a significant 2-way interaction between Valence and Session. Negative relative to neutral lure images, as well as positive relative to neutral lures that were false alarmed failed to differ significantly across Study and Test sessions (negative vs. neutral: z = 0.03, *p* = 0.98, 95% CI [-0.54, 0.56]; positive vs. neutral: z = −0.28, *p* = 0.78, 95% CI [-0.62, 0.47]). Despite the lack of significant interactions, the RE exhibited a significantly different activation for negative relative to neutral false alarmed lures irrespective of session (Main effect of Valence: z = −2.44, *p* = 0.02, 95% CI [-0.87, −0.10]) (Figure 2). No main effects of valence were observed when comparing positive to neutral items (Main effect of Valence: z = 1.73, *p* = 0.08, 95% CI [-0.04, 0.72]). Final exploratory paired samples t-test revealed elevated activation in the RE for negative relative to neutral lure false alarms at Test (*t*(33) = 2.71, *p* = 0.01) and Study (*t*(33) = 2.78, *p* = 0.01). For positive relative to neutral false alarmed lure images, the Study session revealed a similar response (*t*(33) = 2.12, *p* = 0.04), although the Test session did not show significant differences (*t*(33) = 1.57, *p* = 0.13) (Figure 3).

**Figure 2.**
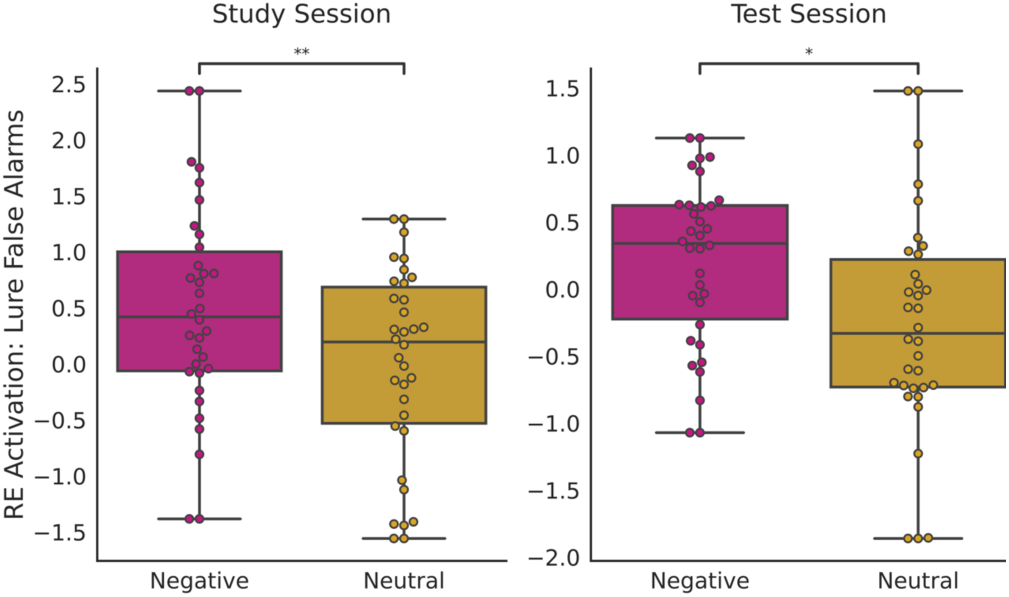
The nucleus reuniens (RE) does not differentially contribute to negative overgeneralization during Study compared to Test sessions (Valence X Session: z = 0.03, *p* = 0.98, 95% CI [-0.54, 0.56]). However, we did observe a significant main effect of valence (Main effect of Valence: z = −2.44, *p* = 0.02, 95% CI [-0.87, −0.10]). Post hoc paired sample t-tests showed that this result was driven by significantly elevated activation in the RE for negative compared to neutral lures that were false alarmed at Test (*t*(33) = 2.71, *p* = 0.01) and Study (*t*(33) = 2.78, *p* = 0.01). n.s. = not significant.

**Figure 3.**
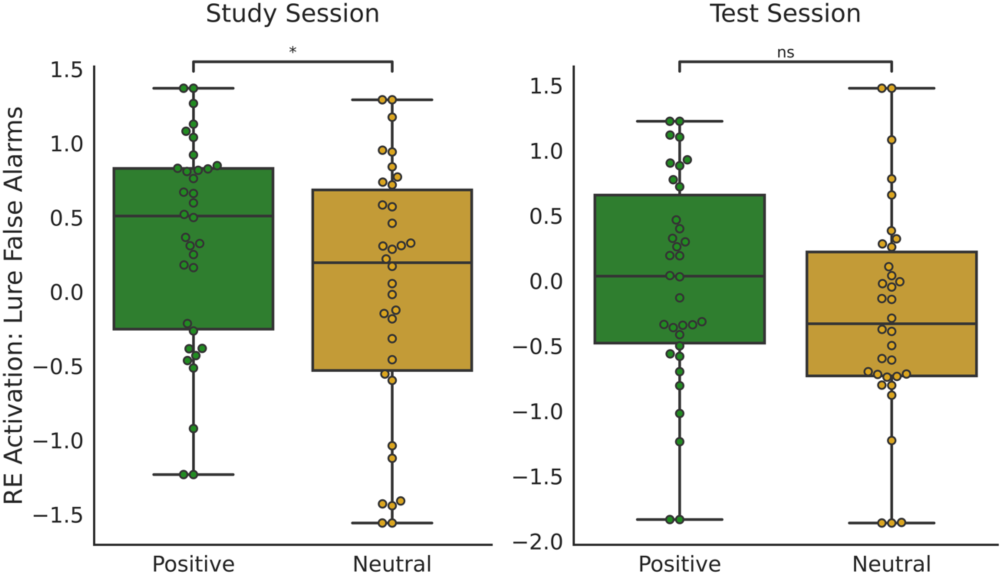
Exploratory analyses showed the nucleus reuniens (RE) contributes to the generalization of positive lure images at Study (*t*(33) = 2.12, *p* = 0.04). No significant differences were observed at the Test session (*t*(33) = 1.57, *p* = 0.13). n.s. = not significant.

### Generalized negative images at Test showed enhanced hippocampal connectivity with the nucleus reuniens

Our next aim was to examine whether the functional connectivity between the RE and CA1 differentially contributed to negative overgeneralization across sessions. The results of our three-way interaction revealed that anxiety severity did not influence functional connectivity differences across Valence and Session (Valence X Session X Pars-6: negative vs. neutral: z = 1.22, *p* = 0.22, 95% CI [- 0.01, 0.04]; positive vs. neutral: z = −1.02, *p* = 0.31, 95% CI [-0.05, 0.02]). Analyses on the interaction between Valence and Session also failed to demonstrate a significant difference in the functional connectivity between the RE and CA1 for negative, as well as positive, relative to neutral lures that were false alarmed across sessions (negative vs. neutral: z = −1.0, *p* = 0.32, 95% CI [-0.30, 0.10]; positive vs. neutral: z = 0.66, *p* = 0.51, 95% CI [-0.16, 0.32]). Further analyses on negative and neutral false alarmed images irrespective of session failed to show significant differences (Main effect of Valence: z = −0.55, *p* = 0.59, 95% CI [-0.18, 0.10]), with positive relative to neutral false alarmed images showing similar results (Main effect of Valence: z = −1.45, *p* = 0.15, 95% CI [-0.29, 0.04]). Final exploratory analyses at Study revealed no differences in the functional coupling between negative and neutral images that were replaced by lures and false alarmed (*Z* = 254, *p* =0.85). Yet, during the Test session, when retrieval is prominent, elevated functional coupling between the RE and CA1 for negative relative to neutral FA images (*t*(31) = 2.63, *p* = 0.01) was identified (Figure 4). Wilcoxon and t-tests exploratory analyses on positive relative to neutral false alarmed images across sessions did not reveal significant differences (Study: *Z* = 264, *p* = 1.0; Test: *t*(31) = 1.42, *p* = 0.17).

**Figure 4.**
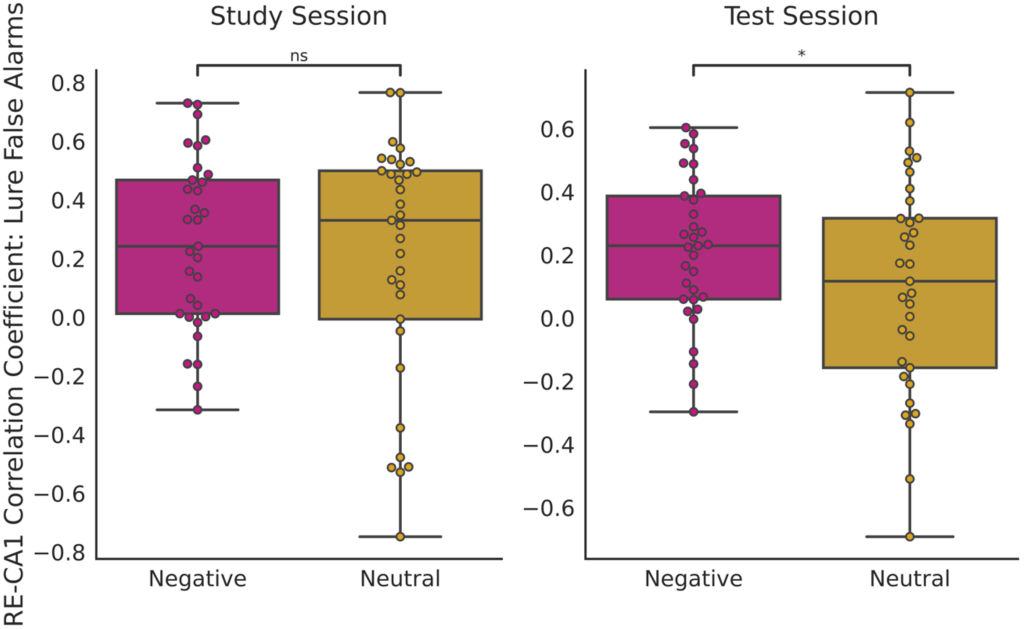
No differences in functional connectivity between the RE and CA1 were evident for negative and neutral lures that were false alarmed across sessions (Valence X Session: z = −1.0, *p* = 0.32, 95% CI [-0.30, 0.10]). Irrespective of session, the RE and CA1 exhibited differences in functional coupling between negative and neutral lures (Main effect of Valence (Main effect of Valence: z = −0.55, *p* = 0.59, 95% CI [-0.18, 0.10]). Post hoc comparisons showed that there was no significant difference in functional connectivity between negative and neutral lures at Study (Z = 254, *p* = 0.85), but at Test the RE and CA1 showed elevated functional coupling for negative relative to neutral lures that were false alarmed (*t*(31) = 2.63, *p* = 0.01).

### Anxiety severity moderates differences in the functional connectivity of the nucleus reuniens and mPFC for neutral relative to negative generalized stimuli at Test

The functional connectivity between the RE and mPFC did not present a 3-way interaction (Valence X Session X Pars-6) across sessions for negative relative to neutral false alarms as a function of anxiety severity (z = - 1.38, *p* = 0.17, 95% CI [-0.05, 0.008]). Subsequent within session analyses, however, revealed a decrease in the functional connectivity for neutral relative to negative generalized images at Test with elevations in anxiety severity (Valence X PARS6: z = −2.65, *p* = 0.008, 95% CI [-0.04, - 0.006]) (Figure 5). The Study session did not show significant differences (Valence X PARS6: z = −0.35, *p* = 0.728, 95% CI [-0.02, 0.017]). Positive relative to neutral stimuli presented similar results, such that no interaction across sessions was observed (z = 0.65, *p* = 0.52, 95% CI [-0.02, 0.04]). Within session analyses showed that neutral relative to positive generalized images at Test were associated with a decrease in the functional coupling between the RE and mPFC as anxiety severity increased (Valence X PARS6: z = 2.26, *p* = 0.02, 95% CI [0.00, 0.04]). Study session showed no significant changes (Valence X PARS6: z = 1.18, *p* = 0.24, 95% CI [-0.01, 0.04]) (see Supplementary section 5 “Anxiety Moderates Differences in the Generalization of Positive Relative to Neutral Stimuli”).

**Figure 5.**
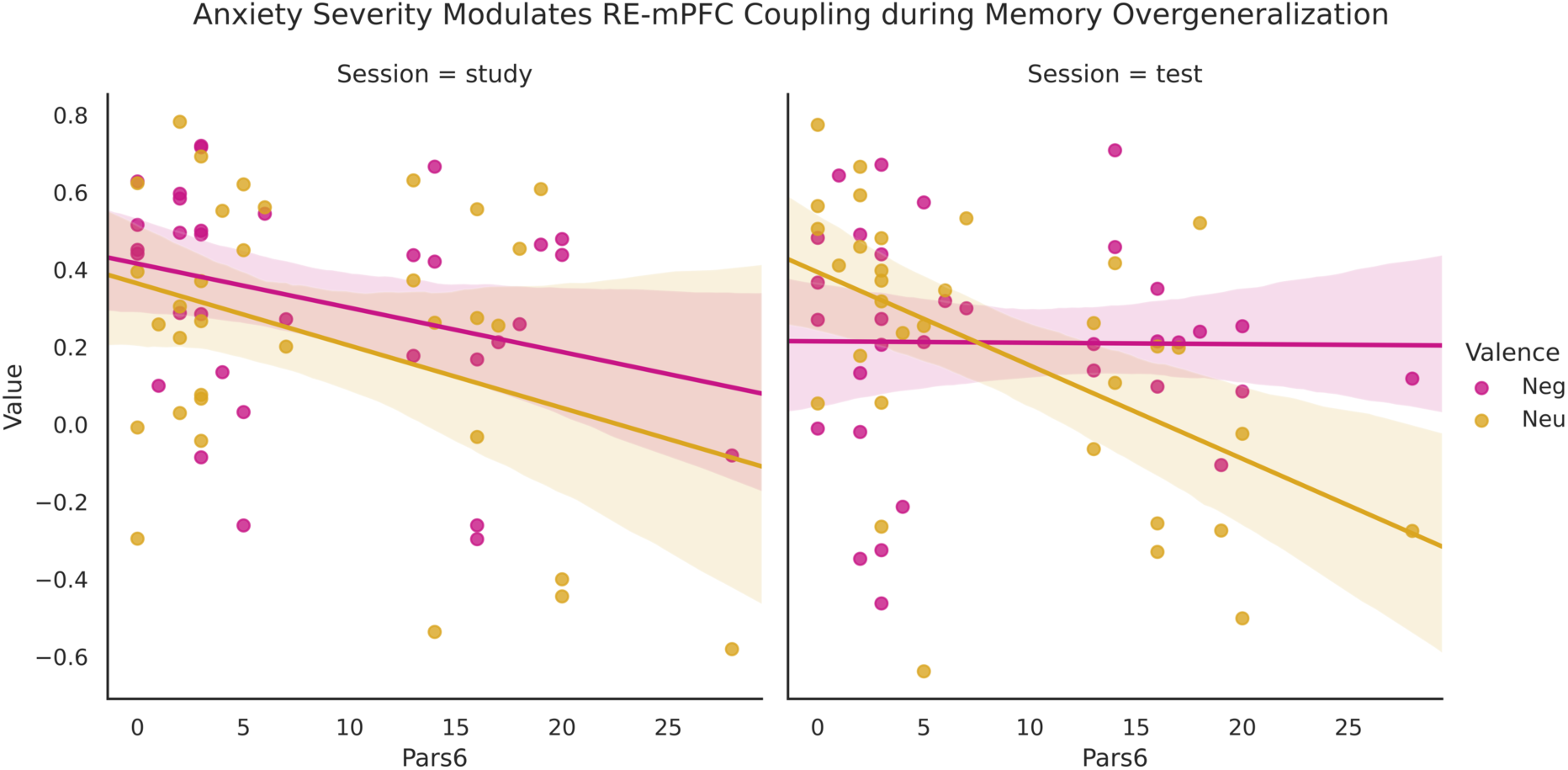
The nucleus reuniens and mPFC did not show significant differences in functional connectivity for negative lures that were generalized across Study and Test sessions (z = 0.61, *p* = 0.54, 95% CI [-0.15, 0.28]). However, at Test, the same regions showed reduced functional coactivation for neutral relative to negative lures that were false alarmed as anxiety severity increased (z = −2.65, p < 0.01, 95% CI [-0.04, −0.01]).

## DISCUSSION

The RE is crucial in the generalization of negative memories. Prior work has focused on the roles of the amygdala, HPC, and the mPFC (8,9,23–25,27,30,67) in memory discrimination and generalization. The RE, however, is well positioned to coordinate and modulate activity in all of these regions (14,32,68,69) and has never been examined in humans in the context of memory generalization. In this study, we observed elevated RE activation at Study and Test sessions for negative relative to neutral lures that were false alarmed, supporting the role of this region in negative overgeneralization both, at encoding and retrieval. We also found that negative overgeneralization was related to the functional interaction between the RE and CA1 during Test and that elevated anxiety severity reduced the RE-mPFC functional coupling at Test for neutral compared to negative false alarmed lures. Our findings support the importance of the RE and that of its frontotemporal functional connectivity in the generalization of aversive and non-aversive memories in youth with differing levels of anxiety.

The RE is instrumental in modulating the specificity of memories during learning. In our study, activations of the RE during the Study session differentiated between negative and neutral items that were subsequently replaced by a lure and false alarmed. These results are consistent with prior rodent studies where changes to the activity of the RE impaired contextual memory discrimination. Reversible muscimol inactivation of the RE during fear learning reduced freezing expression in the conditioned context at test (35). Similarly, optogenetic stimulation of the RE during a contextual fear learning differentially impacted freezing behavior in similar contexts, such that tonic stimulation reduced aversive generalization, whereas phasic simulation enhanced it (32). Our findings support the contribution of the RE in the overgeneralization of negative memories during learning.

The generalization of memories at retrieval is equally dependent on the contribution of the RE. Our data revealed the elevated activation of this region for negative relative to neutral false alarmed lures during the Test session, a finding congruent with rodent models that showing the importance of the RE during recall (32,35). Inactivating the synaptic transmission of the RE 48-hours prior to the generalization test induced a progressive increase in freezing behavior to novel, unconditioned contexts (35). Although one study showed no deficits in aversive memory discrimination following post-conditioning inactivation of RE afferents, the generalization test occurred more than 2-weeks after learning (32). As time continues, additional areas with connectivity to the RE, such as the amygdala (12,70), might work in tandem to improve contextual discrimination. Taken together, our study confirms the role of the RE in aversive memory generalization during recall.

At retrieval, negative overgeneralization is also supported by the functional communication between the RE and CA1. Our results showed the elevated coactivation of the RE with CA1 during the generalization of negative lures at Test. In rodents, electrophysiological stimulation of the rostral two-thirds of the RE elicited excitatory responses in the CA1 (71). In fMRI studies, the DG/CA3 is linked to the storage of distinct inputs as separate elements (pattern separation), while the CA1 is associated with pattern completion, or the reactivation of stored representations by degraded inputs (38–40,72–74). This neurocomputational model has been associated with the overgeneralization of memories (29,40,75,76), even in prepubertal anxious youth (4). Therefore, changes in the activity of the CA1 engendered by the RE could bias the hippocampal network towards an overgeneralization of information. Activity suppression of the RE, for example, was linked to reductions in the activity in the CA1 and enhanced contextual fear generalization (29). The underlying mechanistic basis may be grounded on the capacity of the RE to activate nonpyramidal inhibitory interneurons in the CA1 (71,77) that could likely enhance the firing of unique groups of cells, while silencing the expression of others. Overall, our findings supported the interplay between the RE and CA1 as a neurobiological mechanism of negative overgeneralization.

Impairments in the functional coupling between the mPFC and RE might drive the generalization of non-aversive stimuli in anxiety. Our Test session results revealed reductions in the communication of the mPFC with the RE specifically for false alarmed neutral lures as anxiety severity increased. Previous studies have shown alterations in the activity of the mPFC and RE during the processing of non-aversive stimuli in the context of anxiety (48,78–80). For example, the anterior cingulate cortex displayed hypoactivation during the presentation of neutral facial expressions in patients with anxiety (78–81)and inactivation of the RE enhanced anxiety-related responses (e.g., avoidance) to ambiguous situations (48). The generalization of non-aversive stimuli in patients with anxiety might be driven by alterations in the communication between the mPFC and RE.

Our study had a series of limitations and strengths that warrant attention. Despite utilizing a data-driven approach to define the RE, our region of interest likely included voxels capturing adjacent nuclei, such as the RH. While partial voluming is a common limitation for imaging studies investigating small regions, the RH is connected to cortical and subcortical regions like those of the RE. Further, both nuclei have been shown to be important for learning and memory, as well as anxiety (48,82–84). Yet, our functional understanding of the RE is largely based on rodent models; therefore, its functional contribution to negative overgeneralization is influenced by potential differences in structure across species (70) and the specific tasks used to probe them. For example, whereas most aversive generalization studies are grounded on fear conditioning paradigms, we utilized a task that emphasizes declarative memory processes (i.e., generalization of similar scenes). Despite the limitations, the reported results help elucidate a method for identifying the RE in humans in vivo and its functional contribution to memory generalization and anxiety.

The current study examined the role of the RE and that of its frontotemporal communication in the overgeneralization of negative memories. We confirmed the contribution of the RE by demonstrating its elevated activation and functional connectivity with the CA1 for negative relative to neutral lures that were overgeneralized. Further, elevated anxiety severity reduced the functional communication between the mPFC and RE for neutral generalized items, reflecting impairments in the recollection of non-aversive memory details, potentially driven by poor affect regulation. Overall, we demonstrated for the first time in humans, and in a pediatric sample, the importance of the RE and that of its interaction with the CA1 and mPFC in the overgeneralization of aversive memories, a common feature across anxiety disorders.

## Supporting information

Supplemental Section

## ACKNOWLEDGEMENTS

We would like to thank Drs. Adam Kimbler, Puck Reeders, Nathan Muncy, and TimothyAllen for their continuous help and support throughout this project. The research was supported in part by an R01 from the National Institutes of Mental Health (NIMH) to DLM and ATM (MH116005).

