## Supplemental Section for "Nucleus Reuniens: Modulating Negative Overgeneralization in Periadolescents with Anxiety"

1. **Anatomical Data Preprocessing:** A project specific template was constructed for spatial normalization. As a first step, to minimize errors due to variability in head position across participants, we skullstripped and registered with a rigid-body registration all T1-weighted images to an MNI-152 template using FSL's FLIRT (1,2). Skull-stripped brains roughly in MNI space were used to generate the template using the ANTs buildtemplateparallel.sh script. After template generation, skullstripped brains were warped to this space using ANTs non-linear diffeomorphic warp (ANTs version 2.1.0 (3)). The output warps (combined with the co-registration transformation affines) were utilized to move the RE thalamic mask from template space to individual subject space for extraction of contrast parameter estimates.

2. **Diffusion Imaging Preprocessing.** DWI scans were coregistered to structural scans using the first acquired reference image (b=0) and FSL's BBReg algorithm (4). The parcellation and segmentation file from Freesurfer (i.e., aparc+aseg) was binarized, dilated by 1 voxel, and transformed into DWI space and used as a brain mask. Susceptibility distortions and eddy current corrections were performed on the masked DWI data. Lastly, crossing fibers were modeled within each voxel using FSL's BEDPOSTX (5).

3. **Probabilistic Tractography and K-Means Clustering.** We utilized probabilistic tractographycombined with a data-driven k-means clustering approach to localize the midline limbic thalamicnuclei (Figure S1). Ipsilateral thalamic masks served as seeds for probabilistic tractography (25,000 streamline samples, step length = 0.5, curvature threshold = 0.2, maximum steps = 2000)to 20 ipsilateral cortical and subcortical targets avoiding ventricles. Separate files were created for each cortical and subcortical target, containing the number of random walks that ended at

targets for each voxel within the thalamic masks. The files were vectorized and combined into an "m" x "n" matrix (i.e., 1863 x 20) where "m" was the number of voxels in the left or right thalamus, while "n" was the number of cortical and subcortical targets. We then passed the resulting matrix to a k-means algorithm with a limit of 8-clusters, where voxels served as samples and targets as features implemented in Python (scikit-learn). Each thalamic mask voxel was then assigned a k-means cluster value reflecting their shared connectivity across cortical and subcortical targets and subsequently coerced into 3-dimensional anatomical space. K-means clusters defined by elevated connections to the HPC, parahippocampus, entorhinal cortex, medial orbitofrontal, rostral anterior cingulate cortex, nucleus accumbens, and amygdala, while exhibiting minimal connections to the precentral, postcentral, and paracentral gyrus, as well as the superior frontal gyrus and parietal cortex were binarized. The k-means clusters that were binarized were warped into our study specific template space and averaged together to identify the voxels that overlapped in 80% of our sample. Next, using an established protocol (6) based on the (7) human atlas, we approximated the location of the RE in template space by: 1) identifying the first coronal slice where the interthalamic adhesion was visible (x = 98), 2) calculated the height of the ventral portion of the midline thalamus (i.e., 52% of the entire midline thalamic height – determined by (7)), 3) located an anatomical reference inline with this point – the most dorsomedial segment of the globus pallidus internal, and 4) horizontally segmented the midline thalamus mask (z = 5) into dorsal and ventral portions. All voxels ventral to this division were used as our final RE mask. The resulting midline thalamic and RE masks derived in template space were projected back into each individual subject's functional space to serve as our anatomical regions of interest.

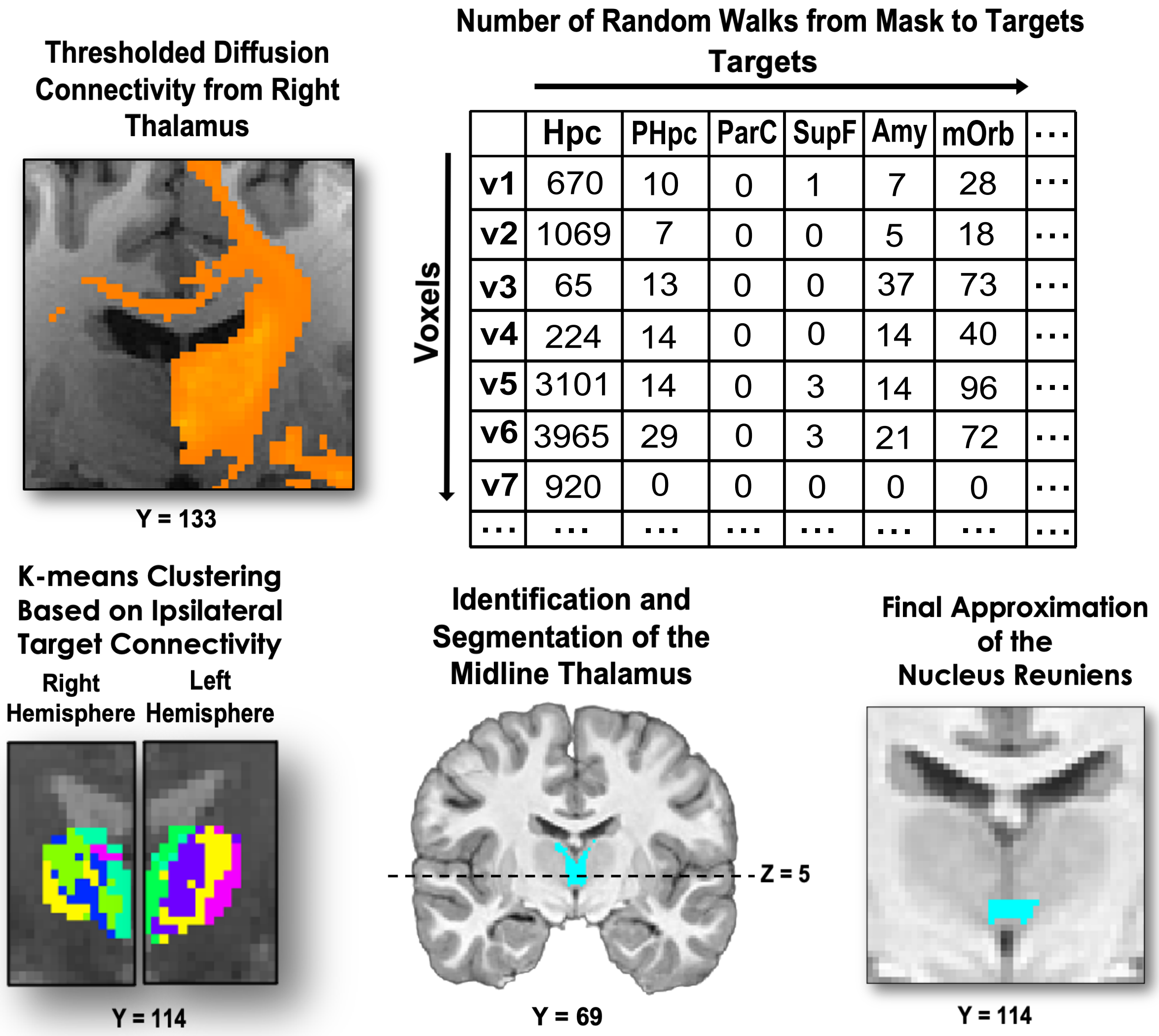


| | Hpc | PHpc | ParC | SupF | Amy | mOrb | ... |
|---|---|---|---|---|---|---|---|
| v1 | 670 | 10 | 0 | 1 | 7 | 28 | ... |
| v2 | 1069 | 7 | 0 | 0 | 5 | 18 | ... |
| v3 | 65 | 13 | 0 | 0 | 37 | 73 | ... |
| v4 | 224 | 14 | 0 | 0 | 14 | 40 | ... |
| v5 | 3101 | 14 | 0 | 3 | 14 | 96 | ... |
| v6 | 3965 | 29 | 0 | 3 | 21 | 72 | ... |
| v7 | 920 | 0 | 0 | 0 | 0 | 0 | ... |
| ... | ... | ... | ... | ... | ... | ... | ... |

4. *The functional coupling between the medial prefrontal cortex and nucleus reuniens did not modulate the generalization of aversive stimuli.* Based on our 3-way interaction results, anxiety did not influence the functional coupling between the RE and mPFC across sessions (z = -1.38, P = 0.17, 95% CI [-0.05, 0.01]). Valence by Session analyses also failed to display significant differences for negative relative to neutral generalized images across sessions (z = 0.61, P = 0.54, 95% CI [-0.15, 0.28]). Analyses on the generalization of negative relative to

neutral lure images did not exhibit a significant difference in the functional connectivity of the mPFC with the RE (Main effect of Valence: z = -1.27, *p* = 0.20, 95% CI [-0.25, 0.05]). Within session analyses showed no significant differences in RE-mPFC functional coupling for negative relative to neutral false alarmed lures (Study: *Z* = 223, *p* = 0.44; Test: *t*(31) = 0.43, *p* = 0.67).

5. **Anxiety Moderates Differences in the Generalization of Positive Relative to Neutral Stimuli**

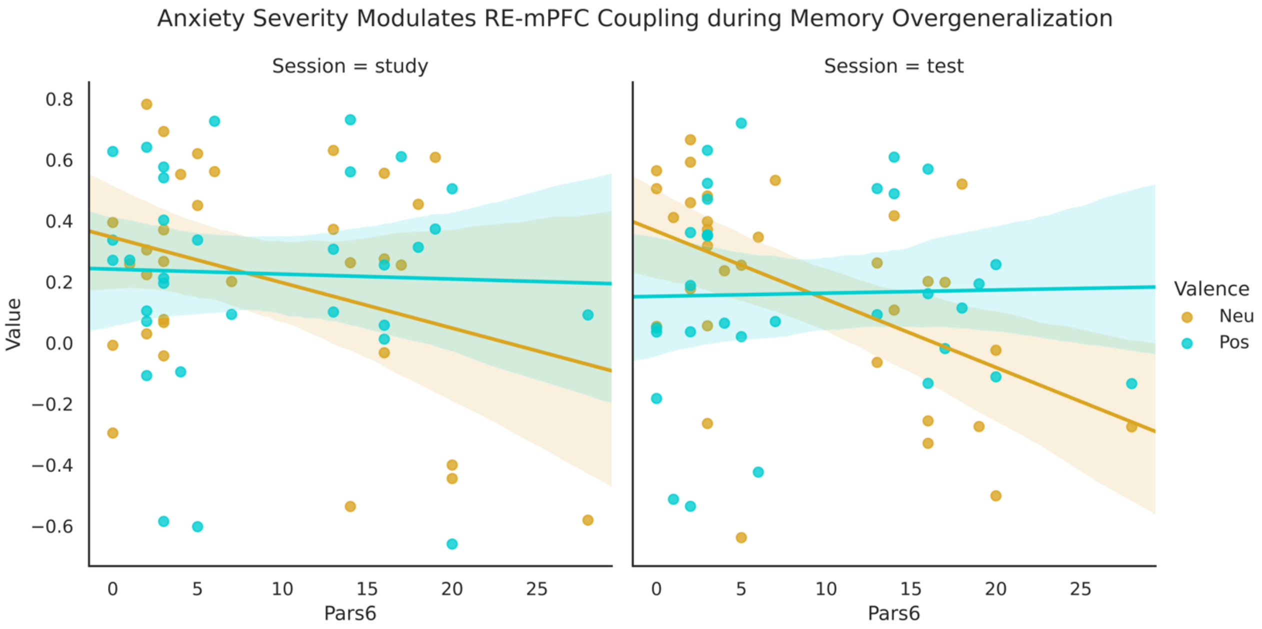
